# Methuselah Proteins in Longevity: Unraveling Their Impact Through Mathematical Genomics

**DOI:** 10.1101/2024.11.03.621698

**Authors:** Sk. Sarif Hassan, Debaleena Nawn, Ankita Ghosh, Moumita Sil, Arunava Goswami, Pallab Basu, Kenneth Lundstrom, Vladimir N. Uversky

## Abstract

This study provides a quantitative and comprehensive analysis of 18 Methuselah (mth) protein variants from fruit flies, focusing on their evolutionary relationships, structural features, and functional roles in aging and longevity. Phylogenetic analysis identified two major clades of mth proteins, with the first clade indicating conserved functions across Drosophila species and the second clade reflecting gene duplication and diversification. The study found five distinct functional subclasses of mth proteins through amino acid frequency and poly-string analyses, linked to their structural diversity and role in longevity. Structural topology and post-translational modifications reveal similarities with G-protein-coupled receptors (GPCRs), suggesting that mth proteins are crucial for signal transduction and cellular health. Variability in propeptide cleavage sites and intrinsic protein disorder further highlight adaptive roles in signaling. The findings underscore the importance of a quantitative and comprehensive approach to studying Methuselah genes, offering insights into their functional versatility and evolutionary dynamics. This enhanced quantitative understanding contributes to advancing research on aging and longevity.

## 1. Introduction

Aging is one of the most complex biological processes, influenced by both genetic and environmental factors [1, 2]. Single-gene mutations that extend lifespan are particularly valuable for understanding the molecular mechanisms underlying aging and longevity determination [1, 3]. Gaining a detailed understanding of the molecular events involved in aging will ultimately help reduce the impact of age-related diseases, improving human health and extending longevity [4, 5].

The fruit fly *Drosophila melanogaster* serves as an excellent model system for dissecting the genetic and cellular basis of crucial biological processes, such as aging [6]. This model allows researchers to uncover parallel mechanisms in vertebrates. The conservation of human disease genes in Drosophila enables functional analysis of orthologs implicated in human aging and age-related diseases [7]. Thus *Drosophila melanogaster* has proven to be one of the most valuable model systems for studying the genetic determination of lifespan [8, 9]. Laboratory studies have demonstrated that its lifespan is highly responsive to genetic manipulations such as induced mutations or artificial selection, with long-lived strains exhibiting up to twice the lifespan of short-lived strains [10, 11]. Despite these findings, critical questions remain, such as whether longevity is primarily governed by many genes of small effect or a few genes of large effect [12, 13, 14]. Furthermore, the identification of aging genes through mutational analyses raises the question of whether these same genes contribute to natural variation in lifespan [15]. Advances in genomic techniques are anticipated to enhance our understanding of the complex genetic architecture of lifespan, particularly in comparing genes that extend lifespan in laboratory settings to those affecting lifespan in natural populations [16].

Single-gene manipulations have led to the identification of candidate aging genes through extended longevity phenotypes, including the Insulin-like Receptor, chico, dFOXO, Indy, and methuselah in the model organism Drosophila melanogaster [15, 17, 18]. Longevity and age-specific mortality patterns are complex traits that vary both within and across species [19]. In model systems like Drosophila, several candidate aging genes have been uncovered through extended longevity mutant phenotypes, such as the G-protein coupled receptor methuselah (mth) [17, 20, 21].

Homozygous individuals with a p-element disruption at mth lived on average 35% longer than the parental strain and showed notable resistance to oxidative stress, starvation, and heat stress [17, 22]. The mth gene encodes a G-protein coupled receptor with seven hydrophobic regions indicative of transmembrane domains and an ectodomain with a ligand-binding site [23]. Lifespan extension is also promoted by disrupting mth ligand activity [20, 24]. Mutations in the stunted gene, which produces two mth peptide ligands, and constitutive expression of antagonist peptide ligands both lead to increased longevity [20, 17, 25]. The mth gene is predicted to encode a protein homologous to several guanosine triphosphate-binding protein-coupled seven-transmembrane domain receptors, suggesting that the organism may utilize signal transduction pathways to modulate stress response and lifespan [26].

However, the mechanism by which mth regulates lifespan remains poorly understood [26]. To gain quantitative insight into mth proteins, a study was conducted that offers a comprehensive analysis of 18 mth protein variants in fruit flies, examining their evolutionary relationships, structural characteristics, and functional roles in relation to longevity.

## 2. Data acquisition

A list of 18 protein sequences of mth from various organisms of Fruit Fly were taken (Table 1). We utilized UniProt, a comprehensive, high-quality, and freely accessible database of protein sequence and functional information, to extract detailed information about each protein of interest [26].

**Table 1:**
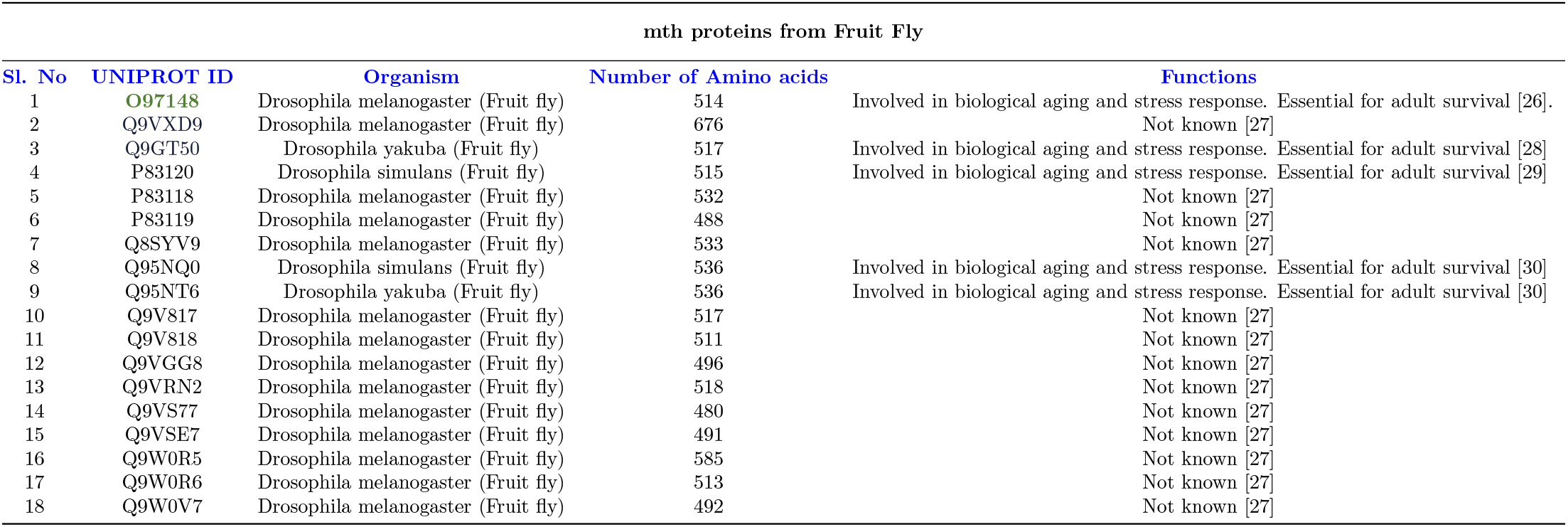
list of 18 mth proteins extracted from UNIPORT database.

**Table 2:** Frequency of homogeneous poly-strings of amino acids across mth proteins.

| mth Proteins | l=1 | l=2 | l=3 | l=4 | l=5 | l=8 | mth Proteins | l=1 | l=2 | l=3 | l=4 | l=5 | l=8 |
| --- | --- | --- | --- | --- | --- | --- | --- | --- | --- | --- | --- | --- | --- |
| O97148 | 450 | 26 | 4 | 0 | 0 | 0 | Q9V817 | 452 | 31 | 1 | 0 | 0 | 0 |
| **Q9VXD9** | 540 | 50 | 8 | 1 | 0 | **1** | Q9V818 | 446 | 27 | 2 | 0 | 1 | 0 |
| P83120 | 456 | 25 | 3 | 0 | 0 | 0 | Q9VGG8 | 410 | 40 | 2 | 0 | 0 | 0 |
| Q9GT50 | 442 | 27 | 4 | 1 | 1 | 0 | Q9VRN2 | 440 | 33 | 4 | 0 | 0 | 0 |
| P83118 | 470 | 28 | 2 | 0 | 0 | 0 | Q9VS77 | 427 | 25 | 1 | 0 | 0 | 0 |
| P83119 | 438 | 22 | 2 | 0 | 0 | 0 | Q9VSE7 | 441 | 22 | 2 | 0 | 0 | 0 |
| Q8SYV9 | 444 | 41 | 1 | 1 | 0 | 0 | Q9W0R5 | 515 | 29 | 4 | 0 | 0 | 0 |
| Q95NQ0 | 474 | 25 | 4 | 0 | 0 | 0 | Q9W0R6 | 442 | 28 | 5 | 0 | 0 | 0 |
| Q95NT6 | 478 | 23 | 4 | 0 | 0 | 0 | Q9W0V7 | 452 | 20 | 0 | 0 | 0 | 0 |

## 3. Methods

Various quantitative features were extracted from 18 mth protein sequences. Methods of feature extractions were discussed as follows.

### 3.1. Sequence homology and phylogeny

Multiple sequence alignment (MSA) was performed using Clustal Omega to identify conserved regions and assess sequence similarity of 18 mth protein sequences[31, 32]. Consequently, a phylogenetic tree was constructed using the Maximum Likelihood method in MEGA X, employing the JTT substitution model, chosen based on the Akaike Information Criterion (AIC) [33]. The tree was visualized using iTOL, highlighting major evolutionary clades [31].

### 3.2. Amino acid frequency distribution across mth proteins

The frequency of amino acids in mth proteins were calculated using *MATLAB* in-build function and the total count of each amino acid was then normalized by the total number of amino acids present in the dataset, providing a relative frequency for each residue. This normalization was necessary to account for differences in sequence lengths [34, 35, 36].

The normalized frequency data were further analyzed to identify patterns and trends in amino acid usage in mth proteins. Statistical measures, including mean, standard deviation, and variance, were calculated for each amino acid’s frequency across the mth proteins. The frequency distribution was visualized using violin swarm-plots, highlighting any amino acids with notably high or low representation. Correlation coefficient was also enumerated for all 18 mth proteins based on frequency percentages of twenty amino acids.

The feature vectors of amino acid frequencies for the mth proteins were used to construct a distance matrix [36]. The Euclidean distance was calculated between each pair of feature vectors to quantify the differences in amino acid composition. This distance matrix provided a measure of dissimilarity between the mth proteins based on their amino acid frequency distributions. A phylogenetic tree was constructed using the distance matrix. The Neighbor-Joining (NJ) method was selected due to its efficiency in handling large datasets and its capability to produce a topology that reflects the input distance matrix accurately [37]. The NJ method was implemented using the MEGA X software, which facilitated the tree construction and visualization [38].

### 3.3. Determining homogeneous poly-string frequency of amino acids in mth sequences

A homogeneous poly-string of length *n* is defined as a sequence consisting of *n* consecutive occurrences of a specific amino acid, as described by Nawn et al. [36]. For instance, in the sequence ‘KKKLLKKLL’, there is one homogeneous poly-string of K with a length of 3, another with a length of 2, and two homogeneous poly-strings of L, each with a length of 2. It is crucial to note that only exclusive and exact occurrences of poly-strings of length *n* are counted.

To determine the maximum length of homogeneous poly-strings for all amino acids across all sequences, we recorded the occurrences of homogeneous poly-strings for all possible lengths, from 1 up to the maximum length, for each amino acid in a given protein sequence [36].

### 3.4. Determining structural topology and domains of methuselah proteins

A comprehensive detailed mapping of the structural topology, domains, and post-translational modifications from the UniPort of each mth protein was extracted [39].

### 3.5. Prediction of proprotein convertase cleavage sites in mth proteins

Cleavage Site Prediction ProP - 1.0 webserver was used to predict the propeptide cleavage sites for arginine (R) and lysine (K) in mth proteins [40]. Default configuration and relevant parameters were used and run the prediction by inputting fasta mth sequences, which will output potential cleavage sites along with associated confidence scores.

### 3.6. Evaluating intrinsic protein disorder of mth protein sequences

Intrinsic protein disorder refers to the lack of a stable, well-defined three-dimensional structure in certain proteins or protein regions under physiological conditions [41, 42, 43]. Proteins that exhibit this characteristic are known as intrinsically disordered proteins (IDPs). Unlike structured proteins, IDPs do not adopt a fixed conformation, remaining flexible even under native conditions [44]. Despite their lack of a stable structure, IDPs are capable of performing diverse biological functions [45].

To evaluate the per-residue disorder propensity in the mth protein sequences, we utilized PONDR® VSL2, a highly accurate standalone disorder predictor [46]. This tool provides per-residue disorder predisposition scores ranging from 0 to 1, where a score of 0 signifies fully ordered residues and a score of 1 denotes fully disordered residues. Residues scoring above the 0.5 threshold are considered *disordered residues*. Those with scores between 0.25 and 0.5 are classified as *highly flexible*, and residues with scores between 0.1 and 0.25 are deemed *moderately flexible* [46].

### 3.7. Determining structural features of mth protein sequences

Protein structural features viz. solvent-accessible surface area (ASA), relative solvent accessibility (RSA), and dihedral angles (*ϕ* and *ψ*), were predicted using NetSurfP-3.0 [47].

## 4. Results and analyses

### 4.1. Sequence homology and associated phylogeny of mth proteins

Sequence homology based phylogenetic relationship among 18 mth proteins reveals proteins Q95NQ0, Q9W0R5, and Q95NT6 (Figure 1) are very much close to each other. This indicates that these proteins are likely orthologs, meaning they are derived from a single ancestral gene present in the last common ancestor of *Drosophila simulans* i.e. DROSI, *Drosophila melanogaster* i.e. DROME and *Drosophila yakuba* i.e. DROYA [48, 49]. The clades containing *Drosophila melanogaster* proteins suggest significant gene duplication and diversification within this species, leading to a variety of protein functions. The phylogenetic tree also indicates that O97148 (DROME) has much similarity with Q9GT50 (DROYA) and P83120 (DROSI). These relationships suggest shared functional and structural characteristics, highlighting the importance of evolutionary conservation in understanding protein function.

**Figure 1:**
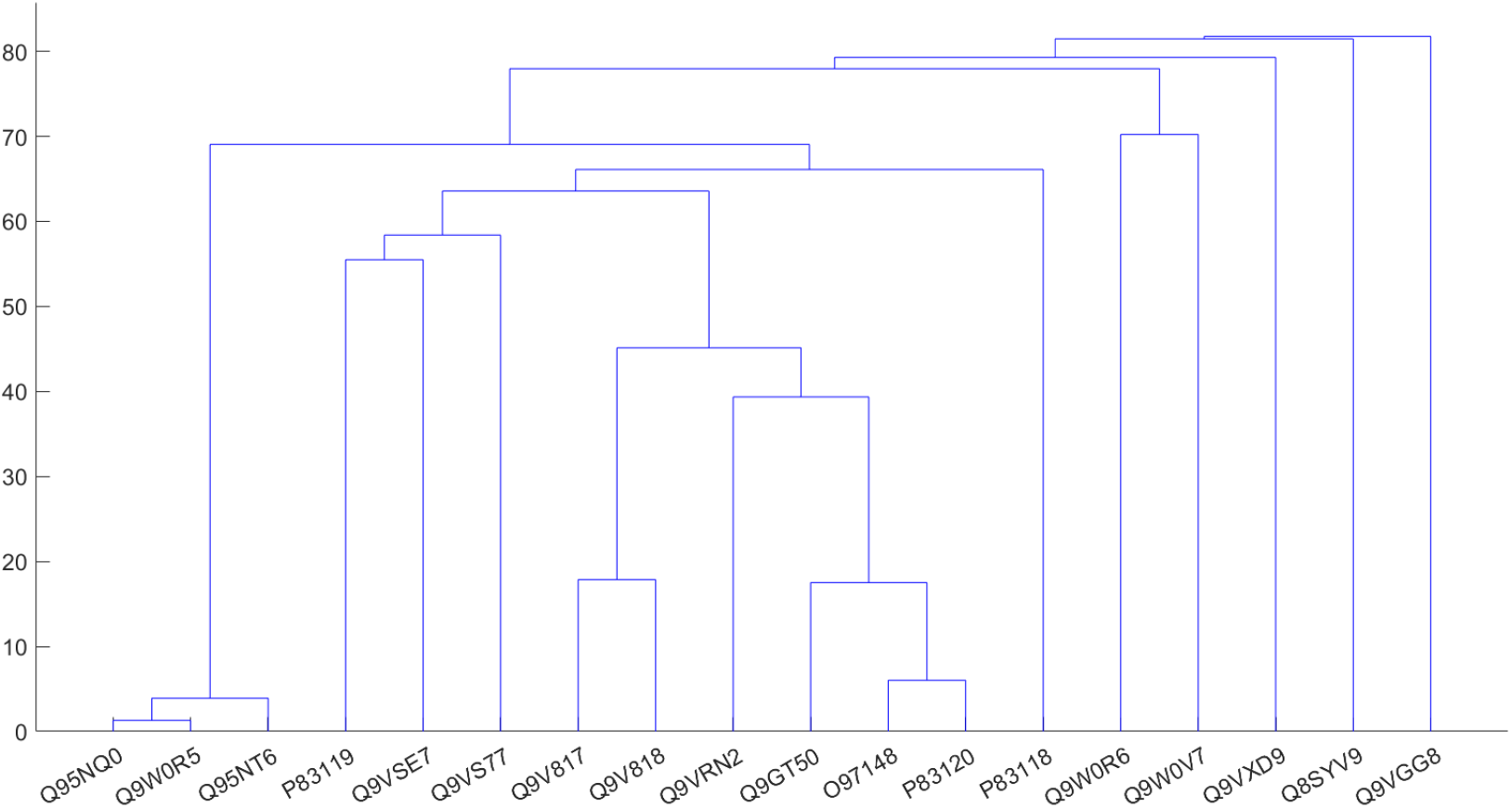
Phylogenetic relationship among mth sequences based on sequence homology

### 4.2. Amino acid frequency distribution across mth proteins

Normalized frequency of each amino acid was calculated across 18 mth proteins (Figure 2. The most abundant amino acid is leucine (L), with an average frequency of 11.12%. High leucine content often correlates with structural roles due to its hydrophobic nature, promoting tight packing in the protein core. Among all amino acids, percentage of leucine is maximum in all mth sequences except Q9VSE7 where isoleucine (I) content is maximum (10.99%). The relatively high presence of serine in most of the mth proteins (average frequency 7.27%) indicates potential roles in enzymatic activity, possibly contributing to biochemical reactions necessary for cellular maintenance and longevity [50, 51]. Among 19 amino acids (excluding leucine), only serine (in Q8SYV9) and isoleucine (in P83119, Q9VS77, Q9VSE7, and Q9W0R6) have normalized frequency greater than 9%. It was observed that hydrophobic amino acid content such as leucine (L), isoleucine (I), valine (V), and phenylalanine (F) was comparatively high (average frequency 11.12%, 7.73%, 6.69%, and 6.59% respectively) and consequently, these proteins likely have a stable hydrophobic core, contributing to their overall structural stability [52]. This stability is crucial for proteins involved in long-term cellular functions, possibly related to the longevity associated with mth proteins [53]. Tryptophan (W) has the lowest average frequency at 2.18%. Despite its low abundance, tryptophan is known for its role in protein-protein interactions and in stabilizing protein structures through hydrophobic interactions [54].

**Figure 2:**
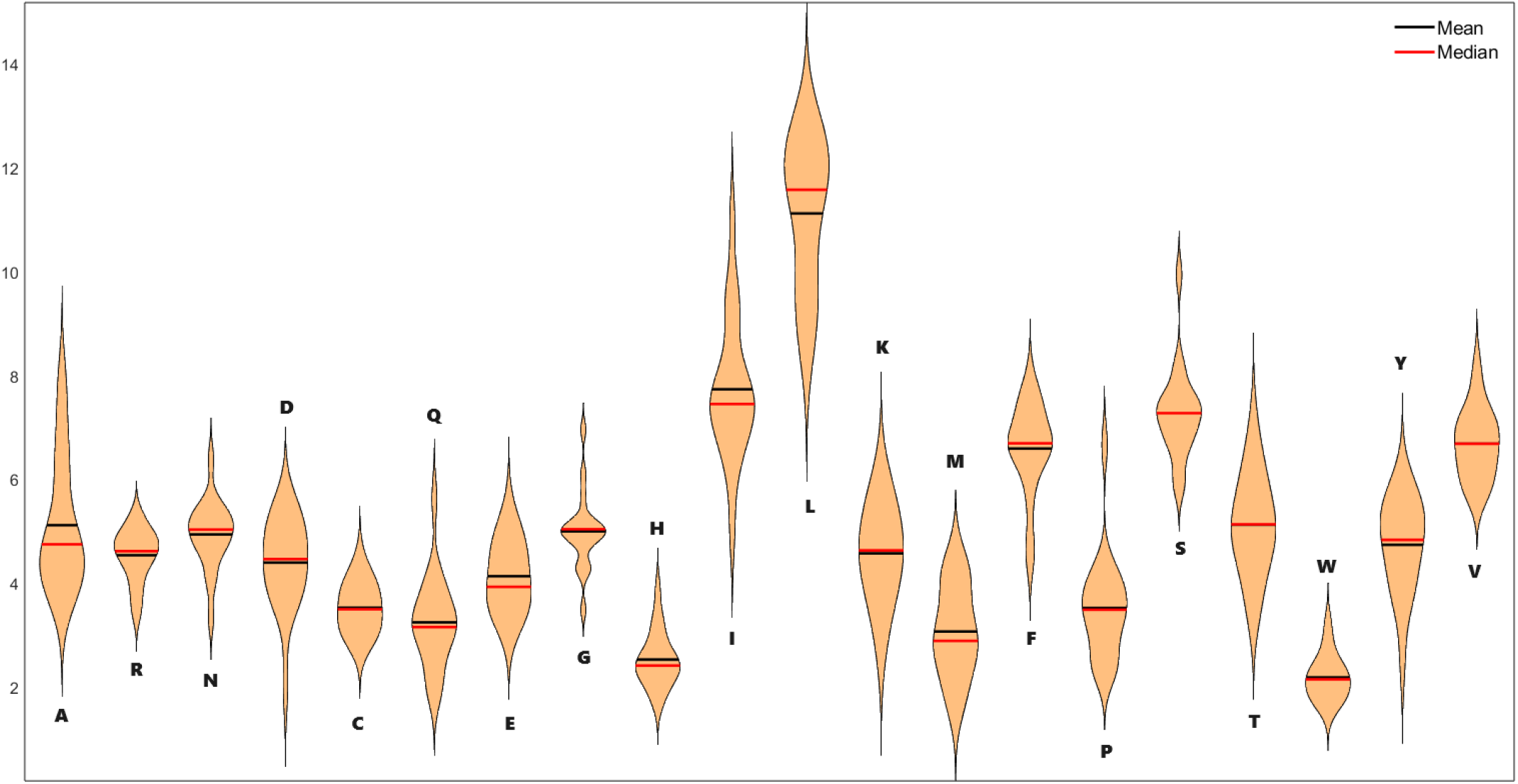
Violin plot of the normalized frequency of each amino acid in 18 mth proteins

Based on the normalized frequency of amino acids in mth proteins, a phylogenetic relationship was drawn (Figure 3). Similar to Figure 1, Q95NQ0 (DROSI), Q9W0R5 (DROME), and Q95NT6 (DROYA) are very close while Q9GT50 (DROYA) formed a cluster with O97148 (DROME) and P83120 (DROSI) based on normalized frequency of amino acids also. However, Q9VGG8 is near to Q9V817 and Q9V818 in Figure 3, but this pattern is not found in Figure 1.

**Figure 3:**
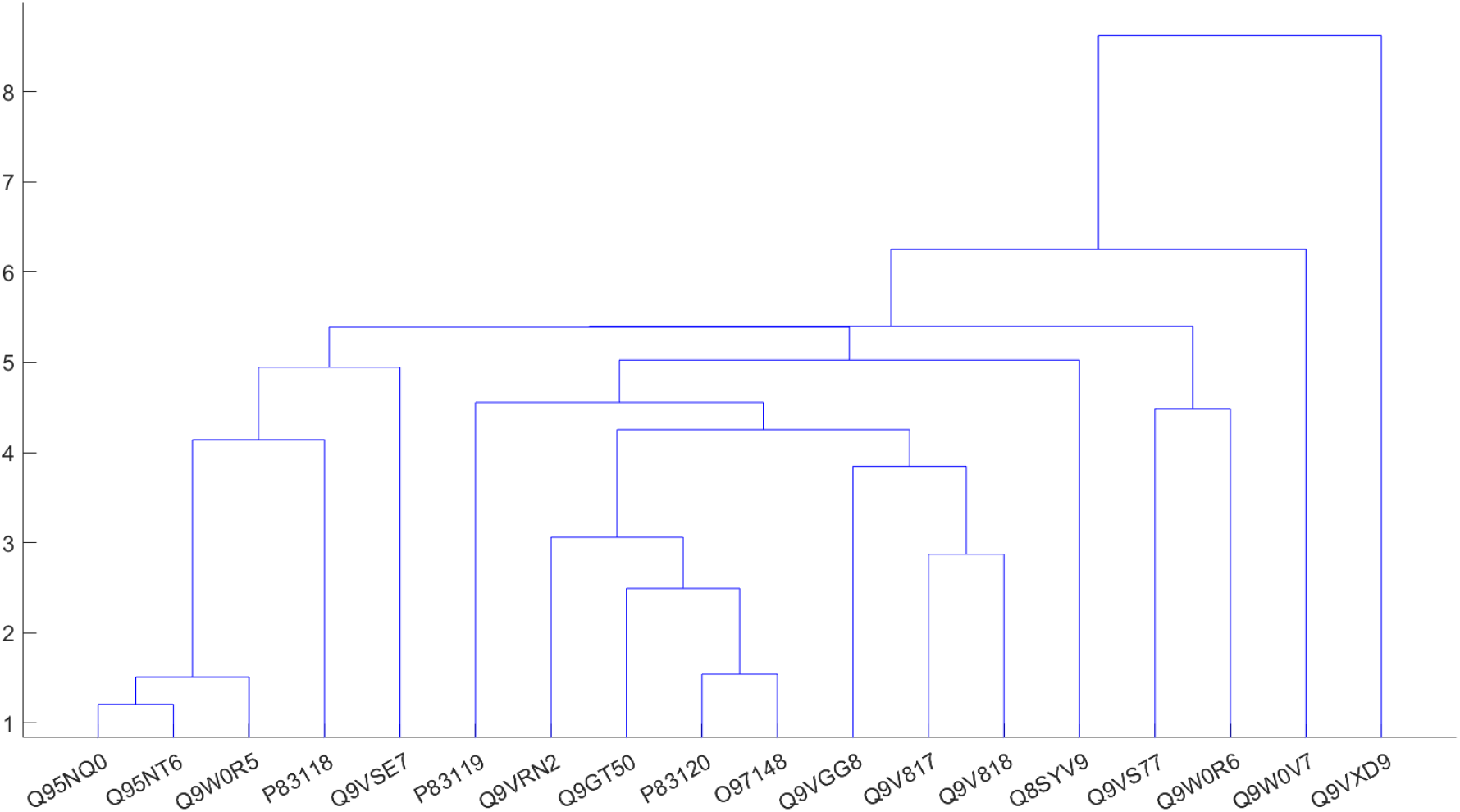
Phylogenetic relationship among 18 mth proteins based on normalized (in %) amino acid frequency distributions

The clustering based on amino acid frequencies hints at structural conservation where certain amino acid compositions are critical for the proteins’ roles in promoting longevity or resistance to stress [55]. This analysis provides a foundation for further experimental validation and practical applications in understanding and leveraging the roles of these proteins in promoting longevity and cellular maintenance [56].

### 4.3. Homogeneous poly-string frequency of amino acids distribution across mth proteins

Table (2) provides the frequency of homogeneous poly-strings of different length *l* in 18 mth proteins. Maximum length of homogeneous poly-strings turn out to be 8. Only Q9VXD9 has one poly-string of length 8 suggesting a unique repetitive structure in this protein and this polystring is composed of Threonine (T). The threonine residues can serve as sites for phosphorylation, impacting the protein’s activity and its involvement in signaling pathways related to aging and stress response [57]. This polystring may also create a unique surface for binding interactions with other proteins or ligands, facilitating the formation of multiprotein complexes crucial for neurogenesis [58]. Additionally, the flexibility provided by the threonine-rich region allows Q9VXD9 to undergo conformational changes in response to environmental cues, enhancing its regulatory capabilities [58]. However, this repetitive sequence could also predispose the protein to misfolding or aggregation, potentially linking it to neurodegenerative processes. Overall, the threonine polystring is likely integral to Q9VXD9’s roles in lifespan regulation and neuronal function [58].

No mth sequence has poly-string of length 6 or 7. Both of Q9GT50 and Q9V818 have single poly-string of length 5 consist of Leucine(L). Q9VXD9, Q9GT50 and Q8SYV9 exhibit single poly-string of length 4 consist of Serine (S), Leucine and Arginine (R) respectively. No mth has poly-string of length 3 composed of Alanine (A), Cysteine (C), Aspartic acid (D), Glycine (G), Histidine(H), Methionine(M), Arginine(R), and Tryptophan (W). All other amino acids have maximum two poly-strings of length 3 in a single mth protein. NNN, PPP, TTT, and YYY (poly-string of Asparagine, Proline, Threonine, and Tyrosine respectively having length 3) was present only in Q9VRN2, Q9VXD9, O97148, and Q9W0R6 respectively. Among them, NNN, TTT, and YYY were found with single occurrence while two PPP was noticed in Q9VXD9.

High frequencies of single amino acids and the presence of longer homogeneous poly-strings in certain proteins suggest specific structural motifs and functions that could be critical for the roles these proteins play in longevity [59, 60].

### 4.4. Structural topology and domains of methuselah proteins

It was noted that most mth proteins possess similar structural topology with signal peptides ranging from 17 to 37 amino acids and consistent chain lengths (Table 3). The presence of topological domains interspersed with transmembrane regions suggests that these proteins span the cell membrane multiple times, typical of G-protein-coupled receptors (GPCRs) [61]. Furthermore, the number and position of transmembrane regions are quite consistent among the proteins, typically showing 7 transmembrane regions. This is characteristic of GPCRs, which are involved in transmitting signals from the outside to the inside of a cell. There is a significant overlap in the domains identified among the mth proteins, indicating conserved regions essential for their function [62]. Several glycosylation sites are conserved among the proteins. Glycosylation is crucial for protein folding, stability, and interactions, indicating that these modifications are important for mth protein functionality [63, 64]. The presence of disulfide bonds across all proteins highlights the importance of structural integrity and proper folding for their function [65].

**Table 3:**
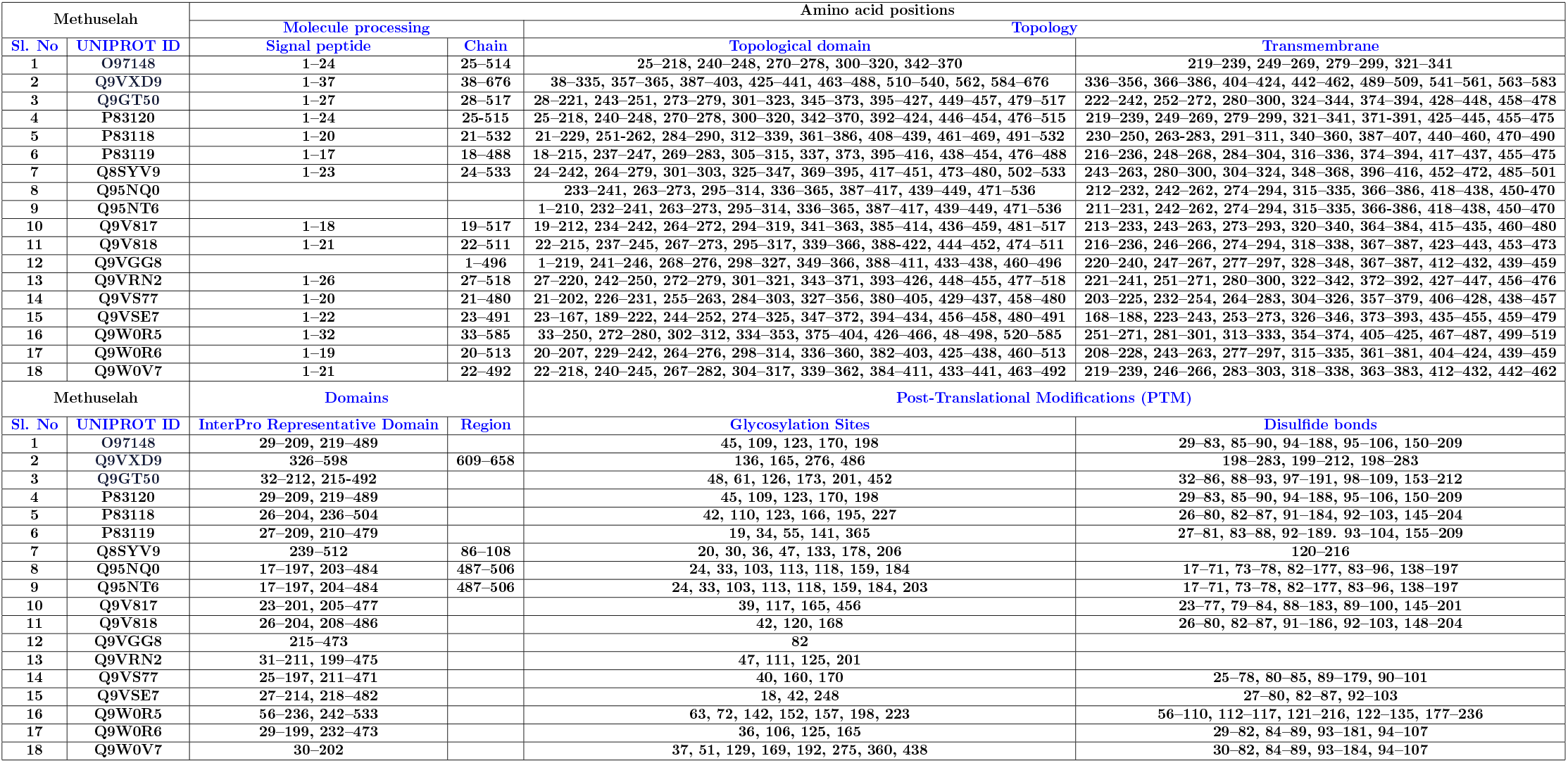
Molecule Processing, Topology, Domains and Post-Translational Modifications (PTM) of mth proteins.

The structural characteristics of mth proteins, resembling GPCRs, suggest they are involved in signal transduction pathways. These pathways are crucial for cellular responses to environmental stimuli, which can influence longevity [66]. Glycosylation and disulfide bonds are important for protein stability and function. Properly folded and stable proteins are more likely to function effectively over time, potentially contributing to cellular health and longevity [67]. The conservation of domains across different mth proteins implies that they might have similar roles in the organism. This functional redundancy could be a protective mechanism, ensuring that essential signaling pathways are maintained, which could positively impact longevity [68]. GPCRs, including mth proteins, are often involved in stress response pathways. Effective stress response mechanisms can help mitigate damage from environmental stressors, thereby promoting longevity.

The mth proteins exhibit highly conserved structural features and modifications that are typical of GPCRs [69, 70]. These characteristics indicate their significant role in signal transduction pathways, which are crucial for maintaining cellular homeostasis and responding to environmental changes. The integrity and functionality provided by glycosylation and disulfide bonds further underscore their importance in cellular processes [71]. Given their roles in stress response and signaling, mth proteins likely contribute to mechanisms that promote longevity, supporting the organism’s ability to cope with environmental challenges and maintain cellular health over time [72].

### 4.5. Predicting arginine and lysine propeptide cleavage sites in mth proteins

Most proteins have a predicted signal peptide cleavage site, suggesting they are likely to be processed and have their signal peptide removed, which is a typical post-transnational modification for secreted and membrane proteins [73]. On the other side it was noticed that two proteins, Q95NQ0 and Q95NT6, do not have predicted signal peptide cleavage sites (‘none’), indicating these proteins may not follow the typical secretory pathway or may lack a signal peptide altogether (Table 4) [74]. The position of the cleavage site varies among proteins, with some occurring early (e.g., P83119 at position 17 and 18) and others much later (e.g., Q9VGG8 at position 43 and 44). The sequences around the cleavage sites also vary, indicating diversity in the signal peptides and possibly in the mechanisms by which these proteins are processed [75].

**Table 4:** Predictions of arginine and lysine pro-peptide cleavage sites in mth proteins.

| Methuselah proteins | Signal peptide cleavage site predicted (between pos) | Pro-peptide cleavage sites predicted |
| --- | --- | --- |
| O97148 | 24 and 25: SYA-DI | Arg(R)/Lys(K): 0 |
| P83118 | 20 and 21: VRS-RD | Arg(R)/Lys(K): 0 |
| P83119 | 17 and 18: TIA-KN | Arg(R)/Lys(K): 1 |
| P83120 | 24 and 25: SYA-DI | Arg(R)/Lys(K): 0 |
| Q8SYV9 | 23 and 24: ASA-QI | Arg(R)/Lys(K): 0 |
| Q95NQ0 | none | Arg(R)/Lys(K): 0 |
| Q95NT6 | none | Arg(R)/Lys(K): 0 |
| Q9GT50 | 27 and 28: TNA-AI | Arg(R)/Lys(K): 0 |
| Q9V817 | 18 and 19: SNA-EI | Arg(R)/Lys(K): 0 |
| Q9V818 | 21 and 22: SNA-EI | Arg(R)/Lys(K): 0 |
| Q9VGG8 | 43 and 44: VTS-HV | Arg(R)/Lys(K): 0 |
| Q9VRN2 | 26 and 27: SSA-EI | Arg(R)/Lys(K): 0 |
| Q9VS77 | 20 and 21: SEA-VI | Arg(R)/Lys(K): 0 |
| Q9VSE7 | 22 and 23: SNA-DI | Arg(R)/Lys(K): 0 |
| Q9VXD9 | 37 and 38: SLA-IE | Arg(R)/Lys(K): 1 |
| Q9W0R5 | 32 and 33: IPG-IP | Arg(R)/Lys(K): 0 |
| Q9W0R6 | 19 and 20: AKS-VE | Arg(R)/Lys(K): 0 |
| Q9W0V7 | 21 and 22: SWG-FH | Arg(R)/Lys(K): 0 |

The majority of the mth proteins listed have “Arg(R)/Lys(K): 0,” indicating no predicted pro-peptide cleavage sites for these amino acids [76]. This suggests that these proteins do not undergo cleavage at these sites during maturation, or such cleavage is not a feature of their post-transnational modification [77]. Only two proteins, P83119 and Q9VXD9, have a value of ‘1’ in the pro-peptide cleavage sites, indicating one predicted cleavage site. This suggests a specific site where enzymatic cleavage could occur during the protein maturation process, possibly affecting protein activation, function, or stability [78].

A prevalent feature among these mth proteins is the presence of a predicted signal peptide cleavage site, which is a hallmark of proteins destined for secretion or localization to specific cellular compartments [79]. The lack of pro-peptide cleavage sites for most proteins suggests that pro-peptide cleavage may not be a common or necessary step in their maturation [80]. Furthermore, the variability in cleavage site positions and sequences indicates functional diversity among these proteins, potentially reflecting different roles or mechanisms of action [81]. The few proteins with predicted pro-peptide cleavage sites may have unique regulatory mechanisms, involving the removal of specific peptide segments to achieve functional maturation [82].

The presence of signal peptide cleavage sites suggests these proteins are likely to be secreted or associated with membranes, playing roles in inter cellular signaling, receptor functions, or other membrane-associated processes [83]. The absence or presence of pro-peptide cleavage sites provides additional insight into the processing and functional regulation of these proteins.

### 4.6. Intrinsic protein disorder of mth proteins

Per residue intrinsic protein disorder of mth proteins were calculated and presented in Figure 4. Intrinsic protein disorder prediction analysis revealed a high variation in the intrinsic disorder in the C-and N-terminals of mth proteins (Figure 4). It also noticed that the highest percentage of moderately flexible residues were present in all mth proteins other than Q9V817, Q8SYV9, Q9V818, and P83118 in whcih highly flexible residues were dominating residues (Figure 5).

**Figure 4:**
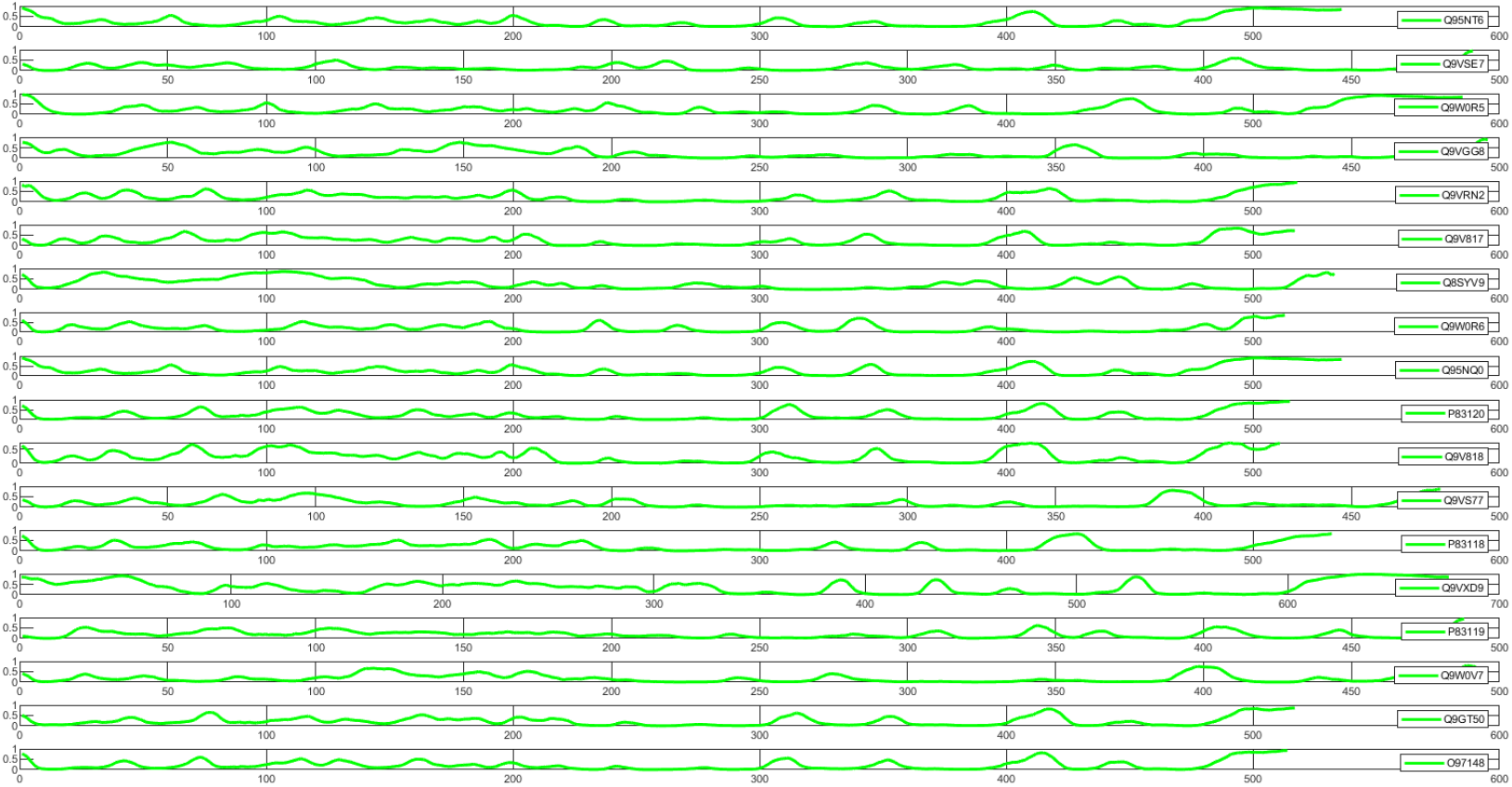
Intrinsic protein disorder residue plots of mth sequences

**Figure 5:**
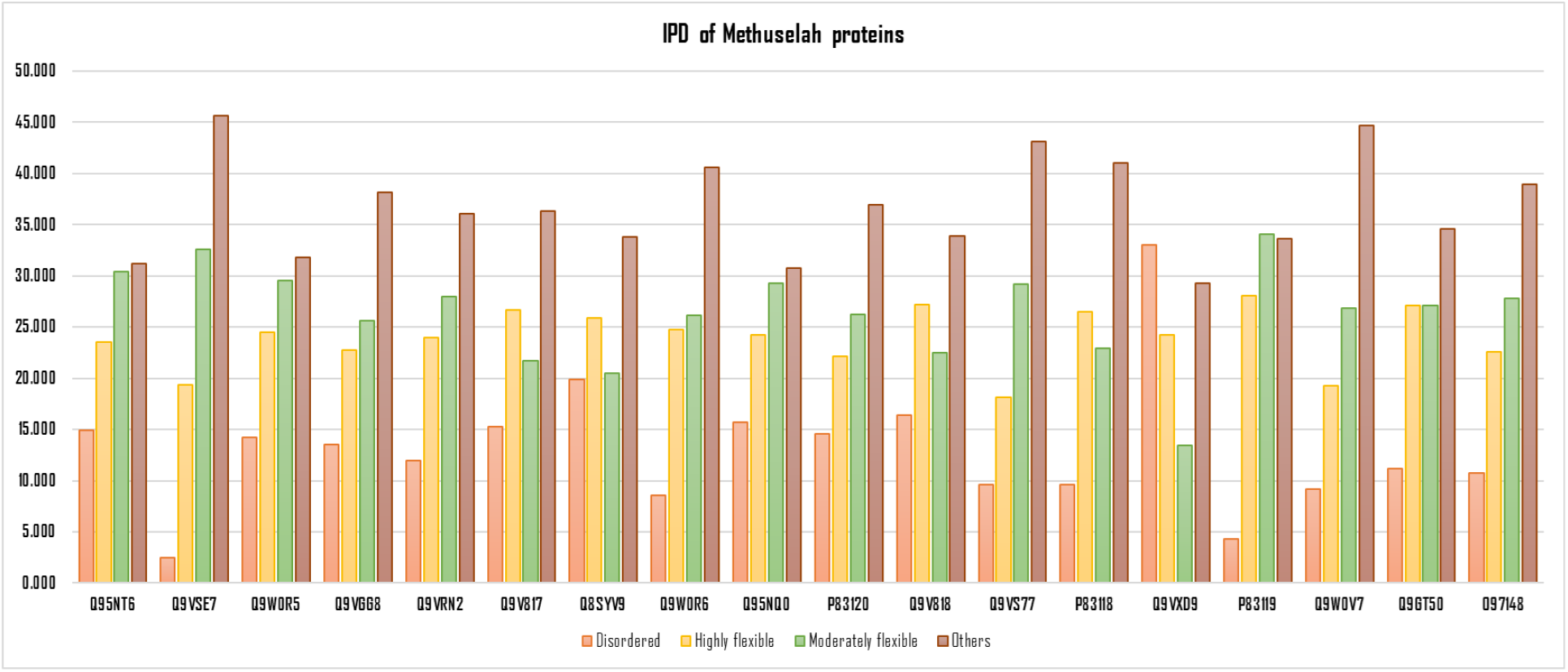
Percentage of intrinsically protein disorder regions of mth sequences

Proteins like Q9VXD9 (32.988%) has significant disordered regions, indicating potential roles in regulation and interactions. Disordered regions are typically involved in signaling and can bind to multiple partners, enhancing functional versatility. P83119 (28.074%) and Q9V817 (26.692%) show high flexibility, suggesting adaptability in their interactions and potential for signaling roles. Flexibility can facilitate binding to diverse molecules, which is crucial for proteins involved in multiple pathways. Furthermore, moderately flexible regions were consistent across proteins, around 20-30%, balancing stability and flexibility, essential for proper function. Almost all mth proteins, except Q9VXD9 and P83119 possessed well-structured regions (‘others’ as mentioned in Figure 5).

The combination of disordered, flexible, and well-structured regions suggests that these proteins are well-adapted to handle various cellular functions [84]. This adaptability is likely important for responding to stress and maintaining cellular health, contributing to longevity [85]. It was noted that the presence of significant disordered and flexible regions suggests these mth proteins can interact dynamically with other molecules, crucial for signaling and regulatory processes [86]. Effective cellular signaling and regulation are vital for longevity, as they help maintain homeostasis and prevent cellular damage [87].

### 4.7. Methuselah protein structural features

For each amino acid in each mth protein, three-state secondary structure viz. relative solvent-accessible area (RSA), and the dihedral angles *ϕ* and *ψ* were computed using NetSurfP-3.0. A snapshot of the predicted structure with RSA and the dihedral angles *ϕ* and *ψ* of the reference mth sequence O97148 was presented (Figure 6).

**Figure 6:**
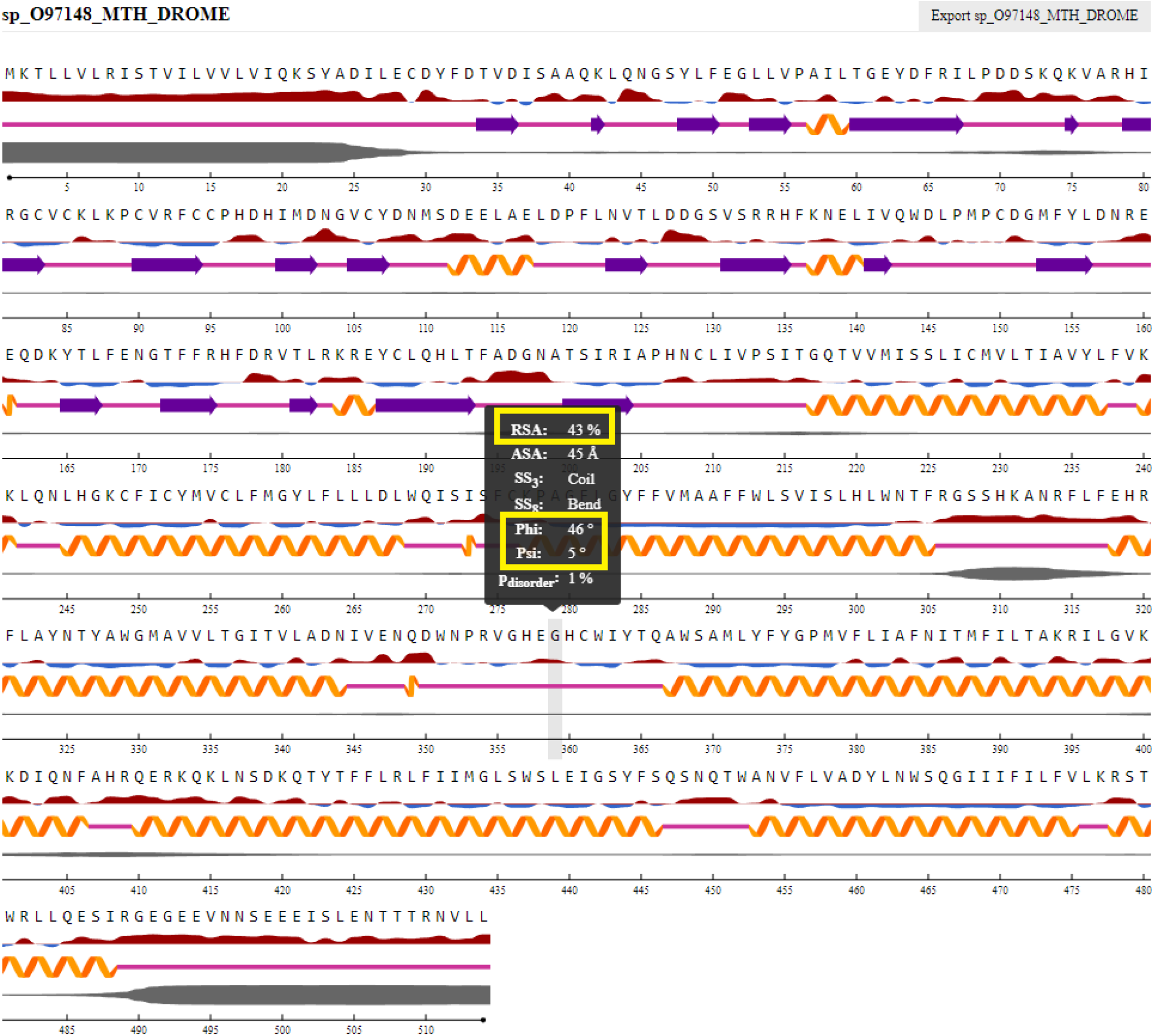
Predicted secondary structural features with RSA and the dihedral angles *ϕ* and *ψ* for the mth protein O97148

**Figure 7:**
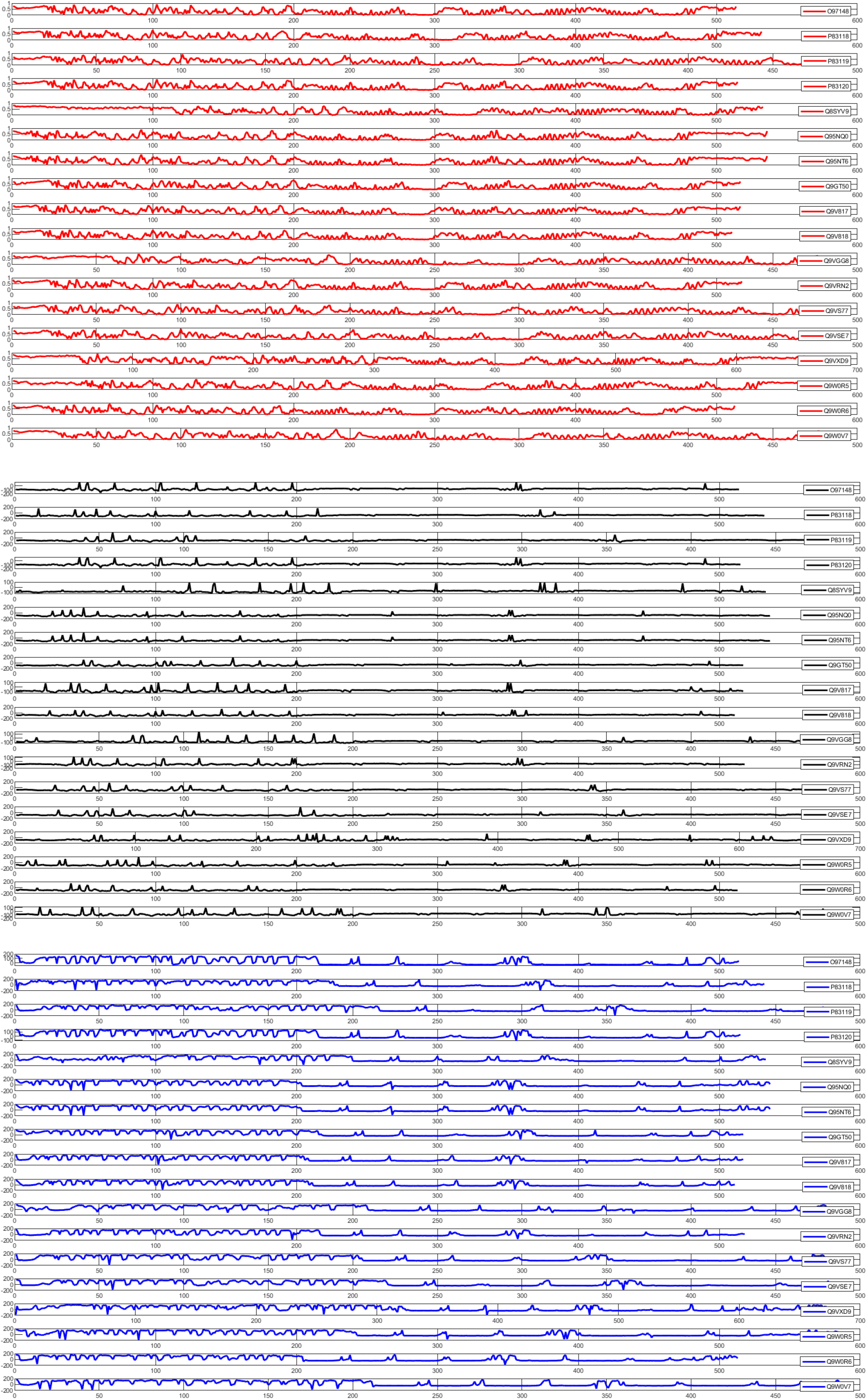
Structural features with RSA and the dihedral angles *ϕ* and *ψ* for all the mth protein sequences

Distribution of RSA, *ϕ* and *ψ* were presented in form of swarm-plots (Figures 8–10).

**Figure 8:**
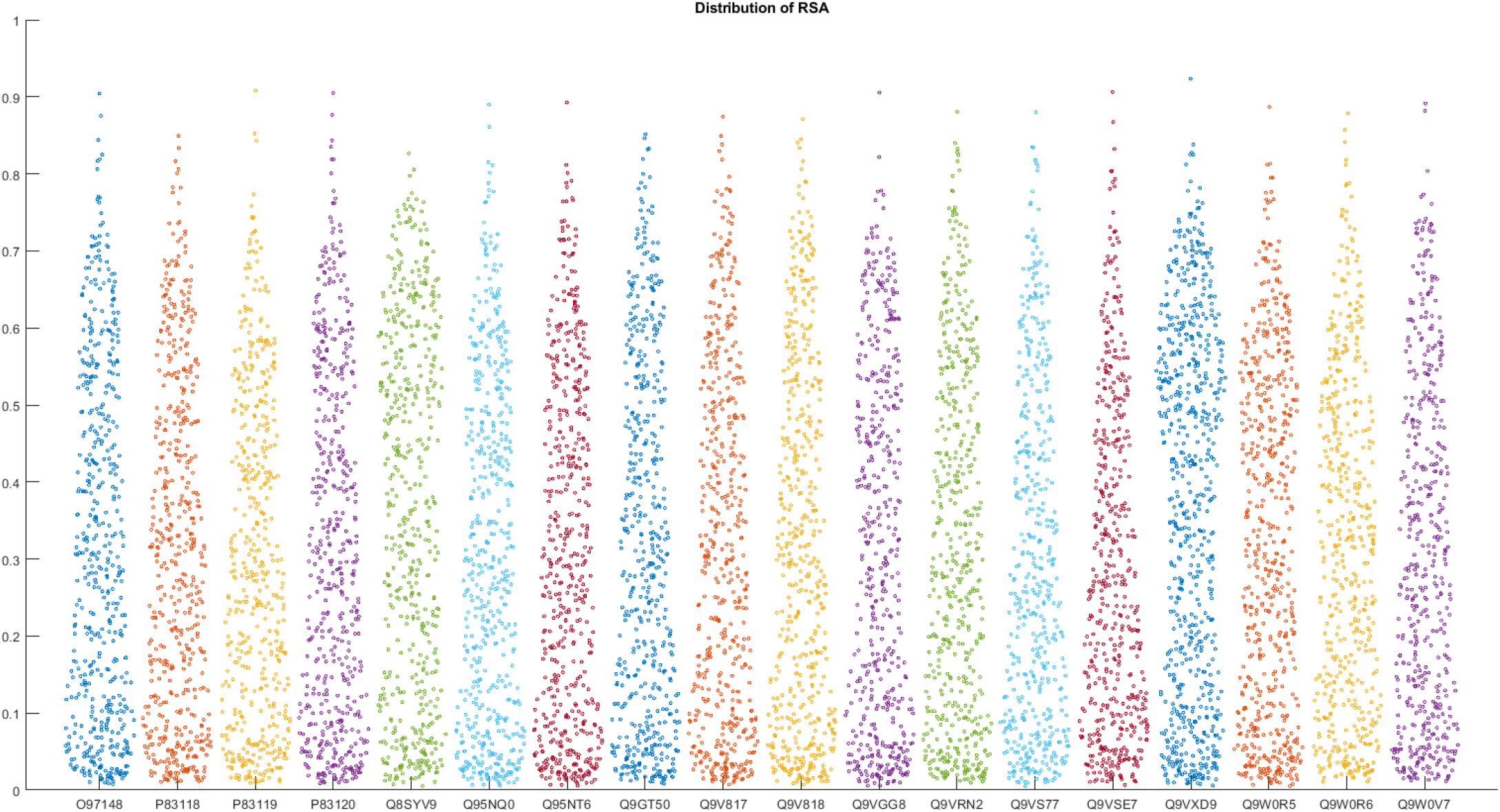
Distribution of RSA

**Figure 9:**
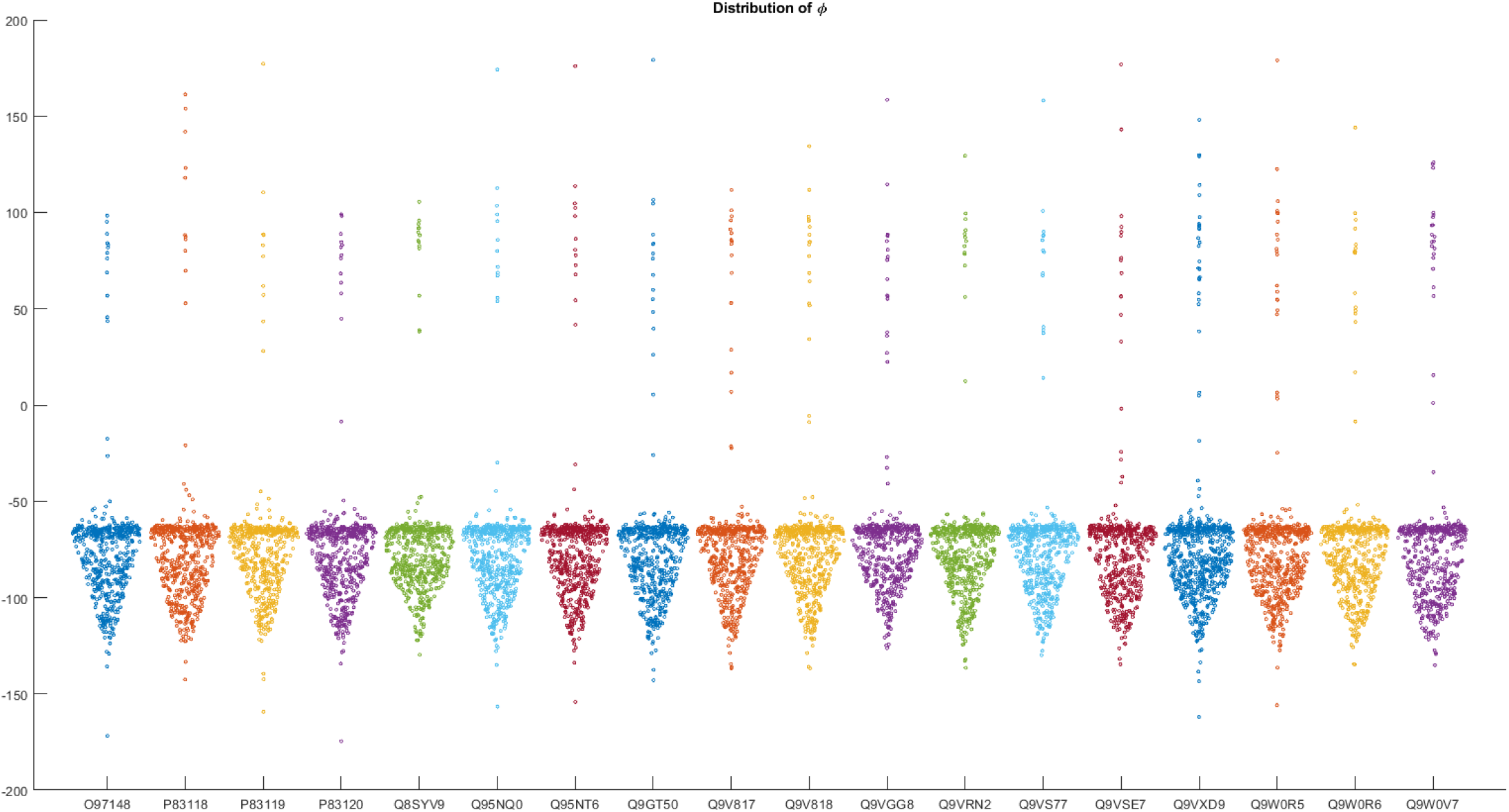
Distribution of *ϕ*

**Figure 10:**
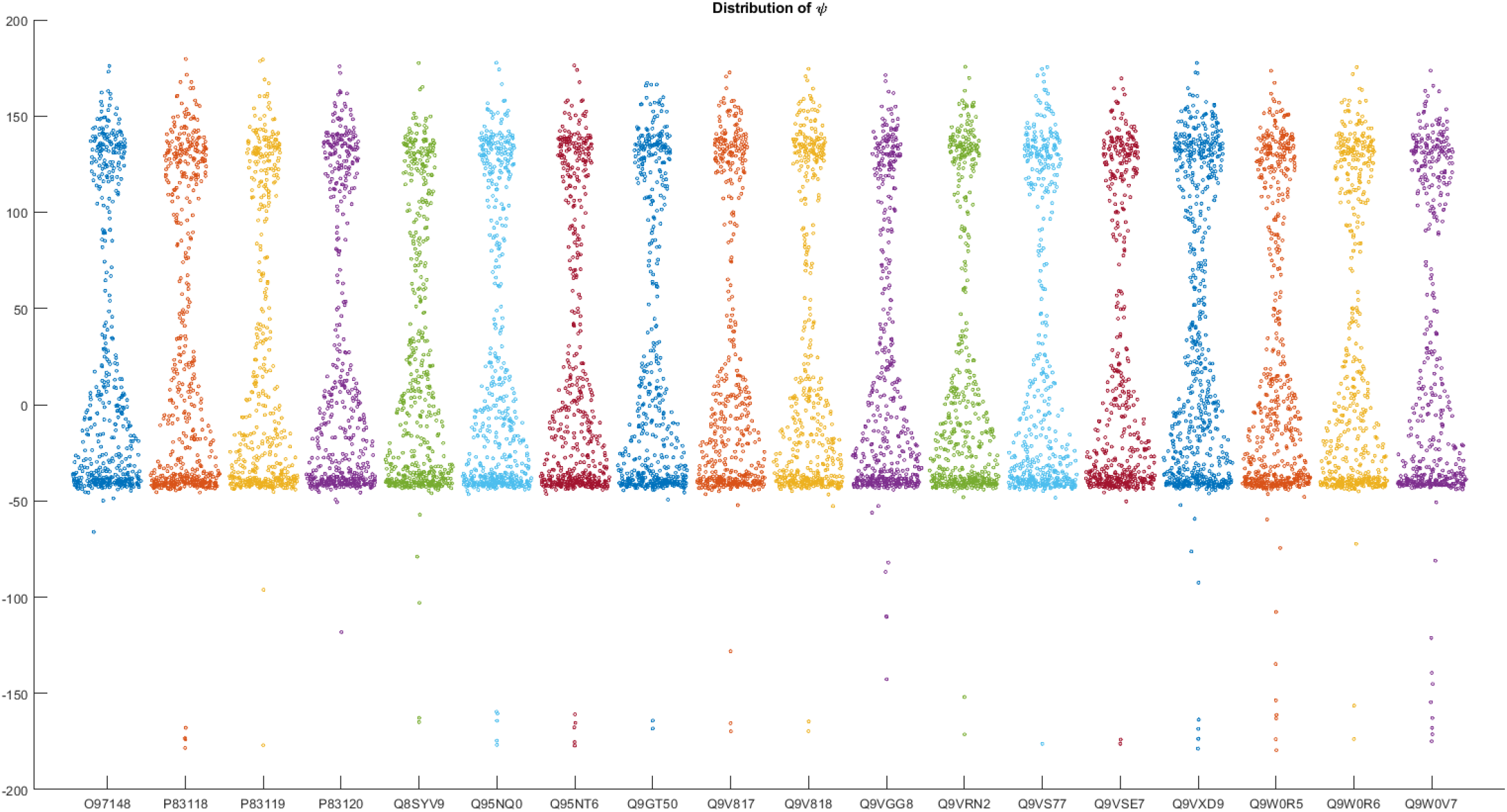
Distribution of *ψ*

Furthermore, mean and standard deviation (SD) of the three features viz. RSA, *ϕ*, and *ψ* for all mth proteins across all amino acid residues were enumerated (Table 5).

**Table 5:** Mean and SD of RSA and *ϕ* and *ψ* for each mth protein.

| Uniprot ID | RSA |  | Phi |  | psi |  | Uniprot ID | RSA |  | Phi |  | psi |  |
| --- | --- | --- | --- | --- | --- | --- | --- | --- | --- | --- | --- | --- | --- |
|  | Mean | SD | Mean | SD | Mean | SD |  | Mean | SD | Mean | SD | Mean | SD |
| *O97148* | 0.32 | 0.23 | -76.91 | 29.56 | 26.05 | 74.03 | *Q9V818* | 0.31 | 0.23 | -74.24 | 33.32 | 23.63 | 75.15 |
| *P83118* | 0.31 | 0.22 | -77.03 | 32.79 | 31.37 | 77.44 | *Q9VGG8* | 0.32 | 0.23 | -73.26 | 30.93 | 19.95 | 71.52 |
| *P83119* | 0.30 | 0.21 | -76.82 | 29.15 | 24.91 | 74.02 | *Q9VRN2* | 0.32 | 0.23 | -75.58 | 30.28 | 20.63 | 73.05 |
| *P83120* | 0.32 | 0.23 | -77.07 | 29.75 | 25.92 | 74.02 | *Q9VS77* | 0.30 | 0.22 | -74.91 | 32.10 | 27.77 | 76.87 |
| *Q8SYV9* | 0.36 | 0.23 | -74.96 | 29.49 | 23.65 | 70.31 | *Q9VSE7* | 0.28 | 0.21 | -76.00 | 32.49 | 24.84 | 74.67 |
| *Q95NQ0* | 0.31 | 0.22 | -76.54 | 30.70 | 24.27 | 75.15 | *Q9VXD9* | 0.37 | 0.23 | -74.34 | 36.02 | 36.69 | 76.11 |
| *Q95NT6* | 0.31 | 0.22 | -76.41 | 30.77 | 24.08 | 75.10 | *Q9W0R5* | 0.32 | 0.22 | -75.61 | 34.34 | 26.54 | 75.74 |
| *Q9GT50* | 0.32 | 0.23 | -76.34 | 31.88 | 27.40 | 75.80 | *Q9W0R6* | 0.32 | 0.21 | -76.20 | 31.49 | 29.51 | 75.77 |
| *Q9V817* | 0.32 | 0.23 | -74.79 | 31.64 | 22.75 | 74.49 | *Q9W0V7* | 0.31 | 0.22 | -74.38 | 37.97 | 28.28 | 79.34 |

Based on this mean and standard deviation of the three features for all 18 mth proteins, a phylogenetic relationship was drawn in Figure 11. From this dendrogram five clusters were developed and they are …

**Figure 11:**
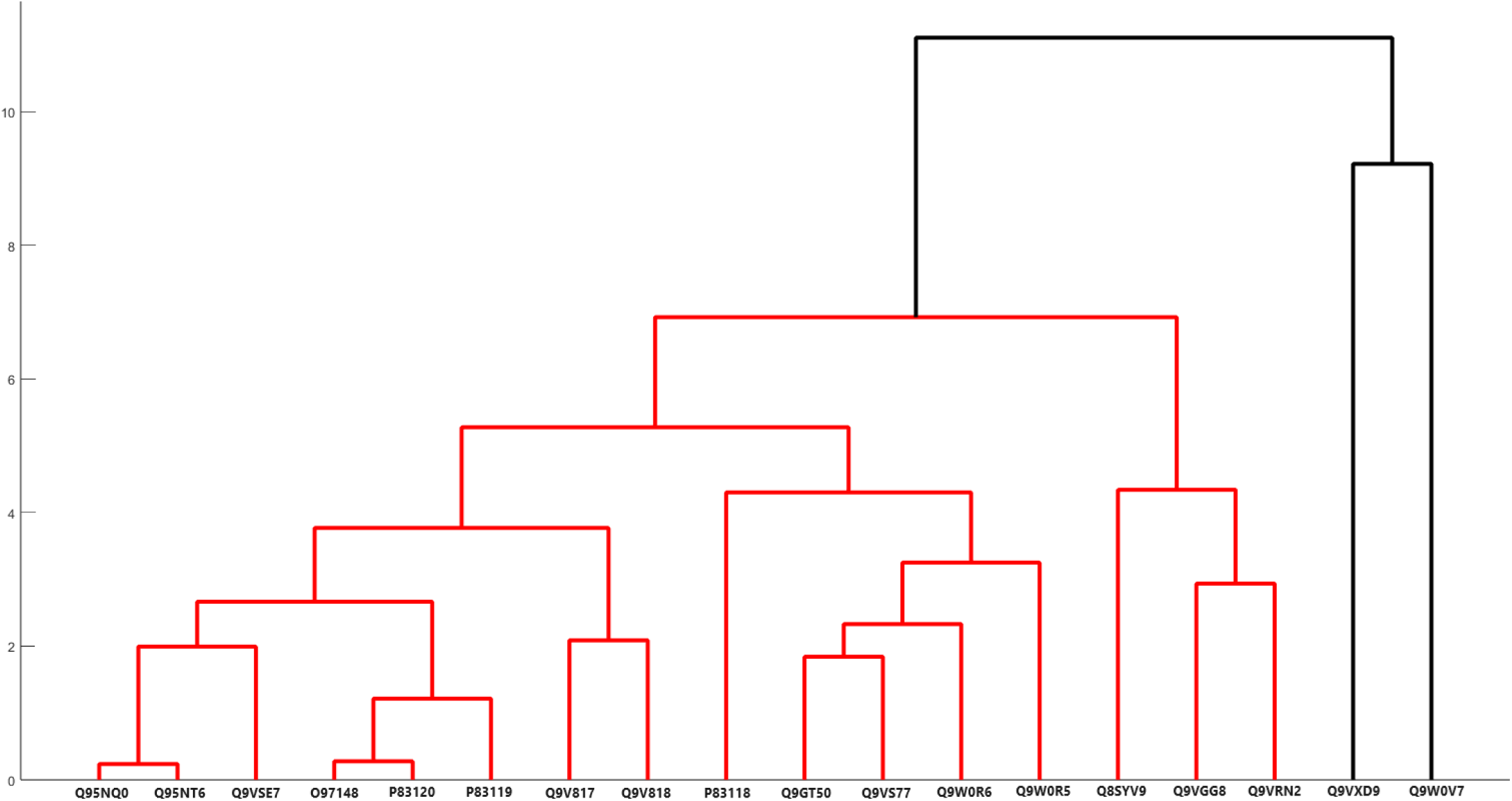
Phylogenetic relationship among mth proteins based on RSA, *ϕ*, and *ψ*.

**Cluster 1:** Q9GT50, O97148, P83119, Q9VSE7, P83120, P83118.

**Cluster 2:** Q95NQ0, Q95NT6, Q9VS77, Q9W0R5, Q8SYV9.

**Cluster 3:** Q9VGG8, Q9VRN2, Q9V818, Q9W0V7.

**Cluster 4:** Q9VXD9, Q9W0R6.

**Cluster 5:** Q9V817.

The mean RSA values range from approximately 0.28 to 0.37. This suggests that the majority of these proteins have moderate solvent accessibility, indicating that they are neither completely buried nor fully exposed. Mean RSA values of the most of the mth proteins hover around 0.31 to 0.32, reflecting a general tendency for these proteins to have similar levels of exposure to the solvent, which could be important for their interactions with other molecules [88]. Furthermore, the moderate RSA means indicate a balance between hydrophobic and hydrophilic regions, which is crucial for protein folding, stability, and function. Proteins with such accessibility are likely to engage in diverse interactions, both hydrophobic and hydrophilic [88, 89]. The distribution of RSA among the 18 proteins indicates a general trend towards moderate accessibility with low variability, suggesting a common structural feature that may be critical for their biological functions [90].

## 5. Discussion and Concluding Remarks

This study provides a quantitative and comprehensive analysis of 18 Methuselah (mth) protein variants from fruit flies, focusing on their evolutionary relationships, structural features, and functional roles related to longevity.

Phylogenetic analysis reveals two major clades among the mth proteins. The first major clade, including Q9W0R5, Q95NT6, and Q95NQ0, suggests these proteins are orthologs derived from a common ancestral gene, indicating conserved functions across species such as *Drosophila melanogaster, Drosophila yakuba*, and *Drosophila simulans* [91]. The second major clade reflects more complex evolutionary dynamics, likely due to gene duplication and subsequent functional divergence. Within *Drosophila melanogaster*, multiple sub-clades suggest extensive gene duplication and diversification, potentially leading to diverse protein functions [92]. The close relationships between proteins such as O97148, Q9GT50, and P36120 highlight conserved functional and structural characteristics [93].

Amino acid frequency analysis identified five distinct clusters of mth proteins, suggesting functional subclasses with varying structural features essential for longevity [94]. Poly-string frequency analysis further emphasizes the structural diversity and functional roles of mth proteins, with notable variations in single amino acid frequencies and repetitive motifs indicating specific functional adaptations [93].

Structural topology and post-translational modifications reveal that mth proteins share a common topology with signal peptides and multiple transmembrane regions, aligning with characteristics of G-protein-coupled receptors (GPCRs). The presence of conserved glycosylation sites and disulfide bonds underscores their roles in protein stability and signal transduction, crucial for maintaining cellular health and promoting longevity.

Analysis of propeptide cleavage sites indicates variability in processing mechanisms among mth proteins, with most proteins predicted to have signal peptide cleavage sites, while some lack these features. This variability suggests diverse functional roles and maturation processes. Intrinsic protein disorder analysis reveals significant variability in flexibility, with some proteins displaying high disorder and others showing more ordered structures. This flexibility is likely important for dynamic interactions and signaling, contributing to cellular health and longevity [95].

The analysis of secondary structure, relative solvent-accessible area (RSA), and dihedral angles *ϕ* and *ψ* revealed three group of mth proteins, reflecting their evolutionary relationships and structural similarities.

The study highlights the balance between evolutionary conservation and diversification among mth proteins, underscoring their significance in longevity. The quantitative and comprehensive approach used provides new insights into their functional versatility and evolutionary dynamics, advancing our understanding of mth proteins in aging and longevity [96].

## Author contributions statement

AG and SSH conceived the problem and theoretical experiments. SSH, DN, VNU, AnG, MS, and PB executed the results and performed the analysis. SSH, DN, VNU, and KL wrote the initial draft. All authors reviewed and edited the manuscript. All the authors checked, reviewed, and approved the final version of the manuscript.

